# Structure of the Thin Filament in Human iPSC-derived Cardiomyocytes and its Response to Heart Disease

**DOI:** 10.1101/2023.10.26.564098

**Authors:** Rahel A. Woldeyes, Masataka Nishiga, Alison S. Vander Roest, Leeya Engel, Prerna Giri, Gabrielle C. Montenegro, Alexander R. Dunn, James A. Spudich, Daniel Bernstein, Michael F. Schmid, Joseph C. Wu, Wah Chiu

**Affiliations:** Department of Bioengineering, Stanford University, Stanford, CA, USA; Stanford Cardiovascular Institute, Stanford University School of Medicine, Stanford, CA, USA; Division of Cardiovascular Medicine, Stanford University School of Medicine, Stanford, CA, USA; Department of Pediatrics, Stanford University School of Medicine, Stanford, CA, USA; Department of Biomedical Engineering, University of Michigan, MI, USA; Faculty of Mechanical Engineering, Technion - Israel Institute of Technology, Haifa, Israel; Department of Chemical Engineering, Stanford University, Stanford, CA, USA; Department of Biochemistry, Stanford University School of Medicine, Stanford, CA, USA; Division of Cryo-EM and Bioimaging, SSRL, SLAC National Accelerator Laboratory, Stanford University, Menlo Park, CA, USA

## Abstract

Cardiovascular diseases are a leading cause of death worldwide, but our understanding of the underlying mechanisms is limited, in part because of the complexity of the cellular machinery that controls the heart muscle contraction cycle. Cryogenic electron tomography (cryo-ET) provides a way to visualize diverse cellular machinery while preserving contextual information like subcellular localization and transient complex formation, but this approach has not been widely applied to the study of heart muscle cells (cardiomyocytes). Here, we deploy an optimized cryo-ET platform that enables cellular-structural biology in human induced pluripotent stem cell–derived cardiomyocytes (hiPSC-CMs). Using this platform, we reconstructed sub-nanometer resolution structures of the human cardiac muscle thin filament, a central component of the contractile machinery. Reconstructing the troponin complex, a regulatory component of the thin filament, from within cells, we identified previously unobserved conformations that highlight the structural flexibility of this regulatory complex. We next measured the impact of chemical and genetic perturbations associated with cardiovascular disease on the structure of troponin. In both cases, we found changes in troponin structure that are consistent with known disease phenotypes—highlighting the value of our approach for dissecting complex disease mechanisms in the cellular context.

## Introduction

Advances in the field of cryogenic electron tomography (cryo-ET) have made it feasible to study macromolecular structure and function in the native environment of cells and at high enough resolution to dissect mechanism^1,2^. These advances have helped overcome technical challenges that have historically slowed the widespread adoption of cryo-ET. For example, limitations with sample preparation focused early seminal work on cell types or peripheral regions that are thin enough to be transparent under transmission electron microscopes^3–6^, but advances like cryo–focused ion beam (cryo-FIB) milling^7,8^ now enable the examination of thicker and more complex cell types^9–13^. Recognizing the potential of these advances to transform study of the complex biology of heart muscle cells (cardiomyocytes), we set out to develop a cryo-ET platform for *in situ* cardiac structural biology.

There is an outstanding need for new approaches in cardiac biology. Heart diseases are the leading cause of death worldwide^14^, and our ability to treat and prevent them is hampered by an incomplete mechanistic understanding of the many different pathways that lead to heart failure and death. While mutations in multiple proteins from the regulatory system of heart muscle contraction have been associated with cardiomyopathies, a mechanistic understanding of how these mutations impact the structure and function of cardiac proteins is rare^15^. Likewise, cardiotoxicity is a major risk factor for patients undergoing cancer treatments, but the mechanisms of cardiotoxicity for different drugs remain unclear^16^. With detailed structural information on the consequences of cardiomyopathy-associated mutations or drug-induced cardiotoxicity, we could accelerate the design of new treatments to combat these diseases^17^. However, the number of components and the complexity of their interactions has limited our ability to reconstitute the system *ex situ*. This has proven a challenge for building models that can connect perturbations like mutations or drug treatment to the function of the larger system.

Cardiomyocytes (CMs) use a complex regulatory system to control our heartbeat^18,19^. Channels found across multiple cellular compartments coordinate to regulate the release of Ca^2+^ into the cytoplasm, which binds to myofibrils to initiate contraction. Myofibrils are macromolecular complexes that directly generate the force of contraction. They are composed of thin and thick filaments interwoven into muscle fibers, with the repeating unit of organization known as the sarcomere. The thick filaments are mostly composed of the protein myosin. The thin filaments are made up of the actin filament, tropomyosin, which follows the actin helical twist, and the troponin complex, which binds both actin and tropomyosin and mediates the initiation of contraction by Ca^2+ 20,21^. When Ca^2+^ binds to the troponin complex, it induces a conformational shift of tropomyosin that exposes myosin binding sites on the thin filament^22,23^. Upon binding to actin, the myosin head mediates a shift in the relative position of the thick and thin filaments to generate the force required for contraction^24–26^. Myofibrils in human cardiomyocytes consist of many additional proteins^27^ not mentioned above that contribute to their proper organization, structure, and function.

An *in situ* structural approach using cryo-ET has the potential to tackle the complexity of cardiovascular structural biology by directly imaging macromolecular components as they are assembled in cells. Indirect and cell-wide effects of drug-induced cardiotoxicity on protein structure and function could be studied using live cells as an experimental system. The impact of cardiomyopathy-associated mutations on protein structure could be studied in a physiologically relevant context that avoids isolation protocols. Initial efforts towards this goal have already been illuminating. Cryo-ET imaging of myofibrils isolated from mouse cardiomyocytes provided insight into the interactions between myosin and actin that drive muscle contraction^28–30^. Imaging of neonatal rat cardiomyocytes with cryo-ET revealed the structure and organization of thin filaments^31^. However, important questions remain about the structure of the myofibril in human cells.

To answer these questions and advance the technology, we deployed a customized cryo-ET platform for studying *in situ* cardiac structural biology in human cells (Fig. 1). First, we focused on human induced pluripotent stem cell–derived cardiomyocytes (hiPSC-CMs) as a model system for cardiac structural biology. We chose hiPSC-CMs because human primary-CMs are readily modified during culturing^32^ and because there are key differences between human and rodent CMs^33^ that limit the applicability of rodent cells for studying human disease. Additional advantages of hiPSC-CMs include the pre-existing availability of hiPSCs derived from several genetic backgrounds associated with cardiovascular disease^34^, CRISPR-based genome editing tools to introduce disease-relevant mutations in isogenic backgrounds^35^, and cell lines with fluorescently tagged proteins that are useful for the study of cardiovascular biology^36^. We designed a cryo-ET platform that was optimized for this model system by incorporating recent advances in sample preparation, data processing, and analysis. For sample preparation, this included micropatterning the electron microscopy grids^37,38^ and using correlated cryogenic light and focused ion beam microscopy within an integrated fluorescent light microscope (iFLM) system. For data processing and analysis, we combined several software packages^39–45^ to maximize the resolution of the reconstructions.

**Figure 1:**
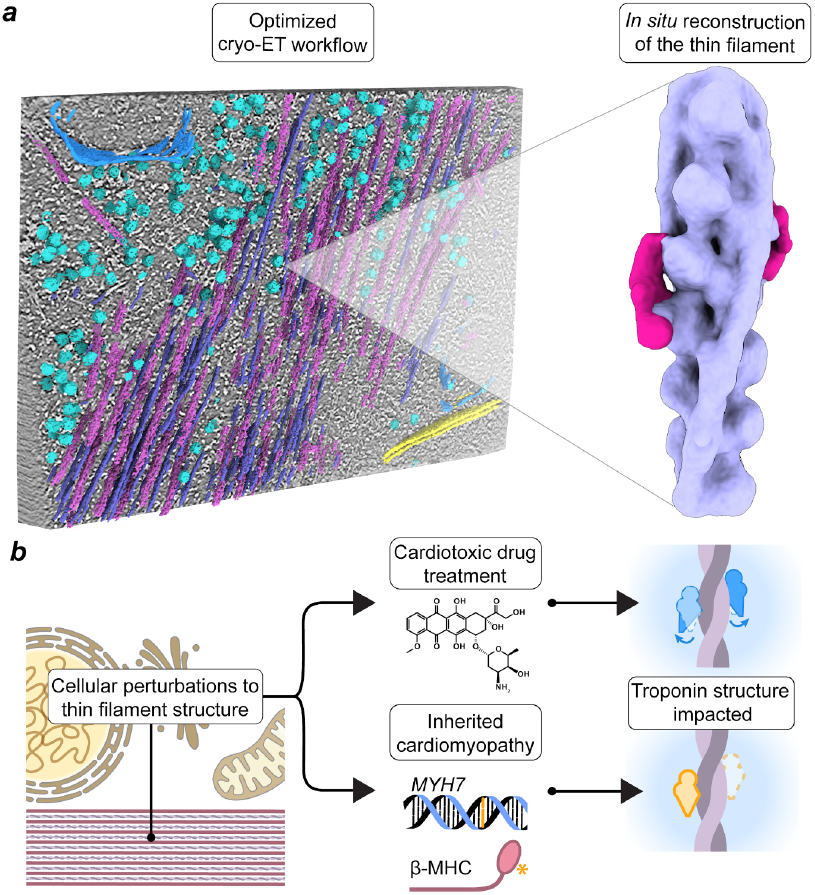
An optimized cryo-ET platform for measuring the effects of cellular perturbations on the structures of cardiac macromole-cules. ***(a)*** (Left) A 3D view of macromolecules segmented from a tomogram of a human induced pluripotent stem cell–derived cardiomyocyte. A slice through the tomogram is shown in black and white. Segmented particles include thick filaments (pink), thin filaments (purple), ribosomes (cyan), vesicles (blue), and mitochondria (yellow). (Right) A map of the thin filament reconstructed from the tomograms. The troponin complex is shown in hot pink while actin and tropomyosin are shown in purple. ***(b)*** Our optimized cryo-ET platform enables new measurements of the impact of cellular perturbations on macromolecular structure. In this work, we studied a cardiotoxic drug (top) and a mutation associated with hypertrophic cardiomyopathy (bottom) and showed that both impact the *in situ* troponin complex structures.

To demonstrate the utility of our platform, we reconstructed the first *in situ* structures of the human thin filament in its cellular context (Fig. 1a). We first reconstructed the core thin filament, consisting of actin and tropomyosin, at sub-nanometer resolution. Next, we reconstructed the troponin complex along with the thin filament. This allowed us to study the impact on thin filament structure from two disease-associated perturbations—treatment with the cardiotoxic drug doxorubicin and a mutation in myosin (*MYH7*^WT/G256E^) associated with hypertrophic cardiomyopathy and hypercontractility^46^ (Fig. 1b). We discovered conformational changes in the structure of troponin consistent with cellular phenotypes previously described in these disease states. This shows the potential of our approach to reveal structural changes that depend on a complex cellular environment, including processes that are indirectly affected by the perturbations introduced. Our approach opens new avenues for dissecting physiologically relevant structural biology in human cardiomyocytes.

## Results

### An optimized cryo-ET platform enables sub-nanometer resolution reconstructions of macromolecules in cardiomyocytes

We started this study by optimizing a cryo-ET platform to visualize macromolecules in cardiomyocytes at sub-nanometer resolution (Fig. S1). We adapted, optimized, and incorporated several advances into the platform to overcome technical challenges associated with cardiomyocytes and achieve sub-nanometer resolution reconstructions.

In the first step of the workflow, we differentiated human induced pluripotent stem cells into cardiomyocytes (hiPSC-CMs) using an established protocol (Methods), then attached them to micropatterned electron microscopy (EM) grids (Fig. S1a). We used micropatterned EM grids to avoid overcrowding of cells, center cells on EM grid squares, and provide spatial cues to improve the organization of contractile proteins within the cells. The pattern consisted of rectangles with a length to width ratio of 7:1, which mimics the dimensions of human adult cardiomyocytes and optimizes contractile function^47^ (Fig. S1c-f). After attaching hiPSC-CMs to the EM grids, we used cryo-focused ion beam (cryo-FIB) milling to prepare approximately 150 nm thick lamellae for cryo-ET.

Two cell lines used in these experiments carried a fluorescent tag fused to α-actinin, a component of the myofibril, at the endogenous locus. For these lines, we used the fluorescent signal from tagged α-actinin to select only hiPSC-CMs with high intracellular organization for cryo-FIB milling and imaging (Fig. S1f). These steps of the workflow benefitted from the use of an Aquilos-2 cryo-FIB/SEM with an integrated iFLM fluorescent microscope, which simplified the steps and increased throughput.

Lamellae from the hiPSC-CMs revealed the characteristic ultrastructures of a cardiomyocyte (Fig. 2a). We detected mitochondria of various shapes and sizes, regions of sarcoplasmic reticulum (SR), and myofibrils that were subdivided into individual sarcomeres at clearly demarcated Z-lines (Fig. 2a). Each of the mentioned ultrastructures play a critical role in maintaining a regular heartbeat in cardiomyocytes.

**Figure 2:**
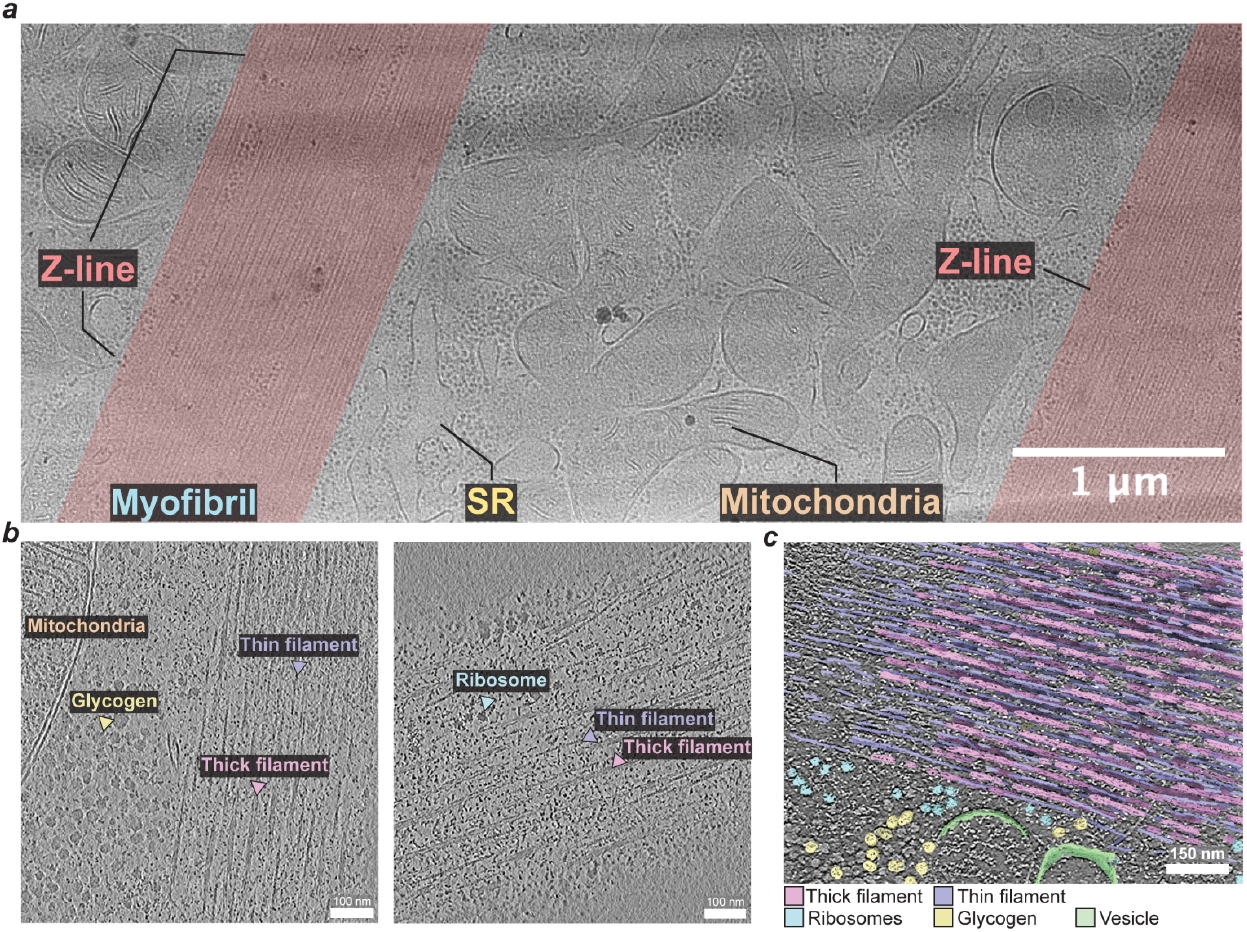
Subcellular arrangements of macromolecules in cardiomyocytes. ***(a)*** Cryo-TEM image collected on a lamella highlighting a region with the myofibrils (highlighted in red), the sarcoplasmic reticulum (SR), and mitochondria. ***(b)*** Z-slices of tomograms from hiPSC-CM lamellae, highlighting the features gly-cogen (yellow), ribosomes (teal), mitochondria (orange), and the thick (pink) and thin (purple) filaments of myofibrils. ***(c)*** Segmentation of a tomogram highlights the variety of macromolecules present, including part of a myofibril with thick (pink) and thin (purple) filaments, vesicles (green), glycogen (yellow) and ribosomes (teal) within the field of view.

We focused on the myofibrils in this work. We collected high-magnification tilt-series images of these regions and reconstructed 3D volumes to better visualize myofibril proteins in their cellular context (Fig. 2b-c). We identified the major components of the myofibril from Z-slices of the 3D volume, including both the thick and thin filaments (Fig. 2b). 3D segmentation of the tomograms allowed us to better visualize how these filaments are found intercalated in the A-band regions of the sarcomere (Fig. 2c) (Movie S1 & S2). In the I-band, the thin filaments extend toward the Z-lines (Fig. 2c). Myosin heads can be detected protruding from the thick filaments, with some bridging to the thin filaments (Movie S1). Many of the tomograms capture other macromolecules and organelles in the immediate vicinity of the myofibril, including ribosomes, glycogen, vesicles, and mitochondria.

To demonstrate the utility of our platform for determining sub-nanometer resolution structures of cardiac macromolecules in the cellular context, we sought to reconstruct the human cardiac thin filament using sub-volume averaging from the tomograms (Fig. S1b, Table S1). We trained a convolutional neural network (CNN) using DeepFinder^42^ to annotate the thin and thick filaments, as well as the membrane and ribosomes in our tomograms. We used the coordinates for the thin filament in untreated control hiPSC-CMs (Fig. 3a) to obtain an 8.7 Å global-resolution reconstruction of the thin filament (Fig. 3b, Fig. S2a). The main components of the thin filaments, filamentous actin and tropomyosin, are clearly resolved (Fig. 3c, Movie S3). In the core of the filament, the resolution ranged from 6–8.5 Å (Fig. 3b). In areas with tropomyosin the resolution ranged from 8–10.5 Å (Fig. 3b), suggesting more flexibility in the position of tropomyosin compared to the actin filament. The helices in individual actin monomers are visible, as expected for the resolution of the map (Fig. 3c, Fig. S3), and coiled-coil models of tropomyosin (PDB-6KN8) can be fit into the density map (Fig. 3c). The overall cross-correlation value from docking the model into the thin filament map is high (0.87).

**Figure 3:**
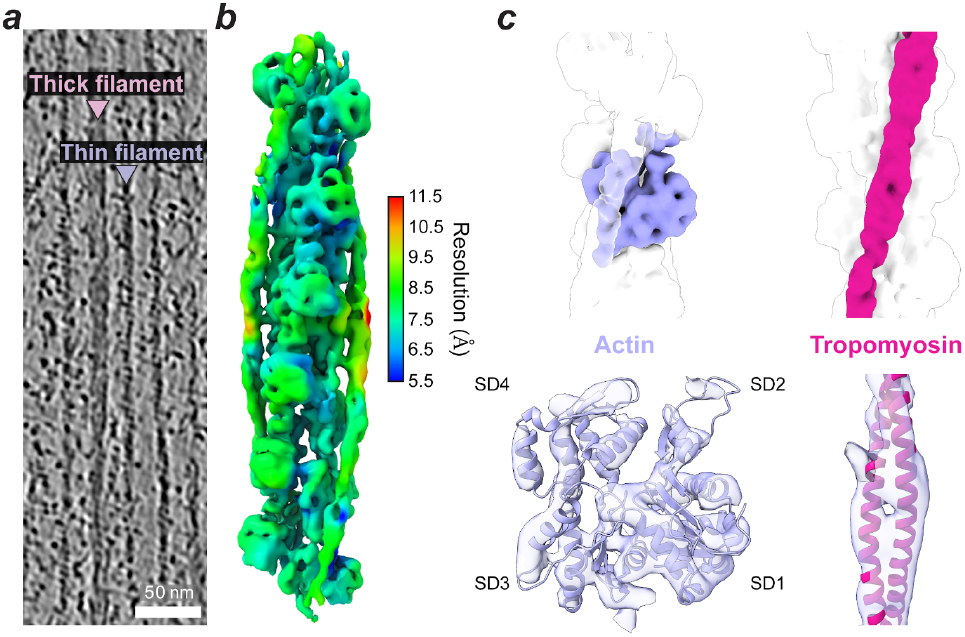
In-cell structures of the human thin filament. ***(a)*** A tomogram slice showing both thick (pink) and thin (purple) filaments. ***(b)*** The structure of the repeating unit of the thin filament, reconstructed from hiPSC-CMs. Actin and tropomyosin subunits are both visible. The resolution of the reconstruction is represented as a colormap. ***(c)*** A molecular model of the thin filament was fit within the *in situ* reconstruction. (Top left) The position of a single actin subunit (purple) is highlighted within the larger reconstruction (transparent white). (Bottom left) A molecular model of actin (PDB: 6KN8) fits within the *in situ* map. Individual helices can be resolved within the actin subunit. The 4 subdomains (SD1-4) are labeled. (Top right) Tropo-myosin (hot pink) is highlighted within the larger reconstruction (transparent white). (Bottom right) A molecular model of tropomyosin (PDB: 6KN8) fits within the *in situ* map.

The current model of the cardiac contraction cycle^48,49^ posits that tropomyosin shifts relative to actin in response to Ca^2+^ binding in order to expose myosin binding sites on the actin. In this model, sarcomeres are made up of a mixture^50^ of three conformational states of tropomyosin: the low Ca^2+^ B-state, the high Ca^2+^ C-state, and the myosin-bound M-state^51–53^. Ca^2+^ and myosin binding to the thin filament increases the relative populations of the C-state and M-state, respectively.

To determine the functional state of our *in situ* map, we compared it to previously determined maps of the B-state (EMD-0728), C-state (EMD-0729), and M-state (EMD-13996) from reconstituted thin filaments^54^ (B and C) and isolated myofibrils^29^ (M). The position of tropomyosin relative to actin in our map is in between the C- and M-state (Fig. S4c). This is consistent with thin filaments in the C-state being a major contributor to the reconstruction.

To our knowledge, this is the first sub-nanometer resolution reconstruction of the thin filament in its cellular environment. This result demonstrates that our platform can achieve reconstructions at a high enough resolution to track functionally important changes in human cardiac proteins, as they occur inside cells. Tracking structural changes in the sarcomere *in situ* will enable new mechanistic insights into complex phenomena that cannot be reconstituted outside of the cell. To test this potential application, we expanded our analysis to include the troponin complex, a regulatory complex of the thin filament, and determined the impact of two perturbations associated with cardiovascular disease on thin filament structure.

### The troponin complex adopts a variety of conformations in response to cellular perturbations

Reconstructing the structure of thin filaments in hiPSC-CMs gave us an opportunity to explore their local neighborhoods within the tomograms, identify additional binding partners, and expand our model of the sarcomeric proteins. Using the thin filament reconstructions, we focused on regions at the filament crosspoints where the troponin complexes are located (Fig. 4a). Classifying and refining these regions of interest, we reconstructed the structure of the troponin complex bound to the thin filament (Fig. 4b). Focusing first on the reconstruction from control experiments in unperturbed hiPSC-CMs, our reconstruction has a global resolution of 18 Å (Fig. S2b) and contains the core components of both the upper and lower troponin complexes.

**Figure 4:**
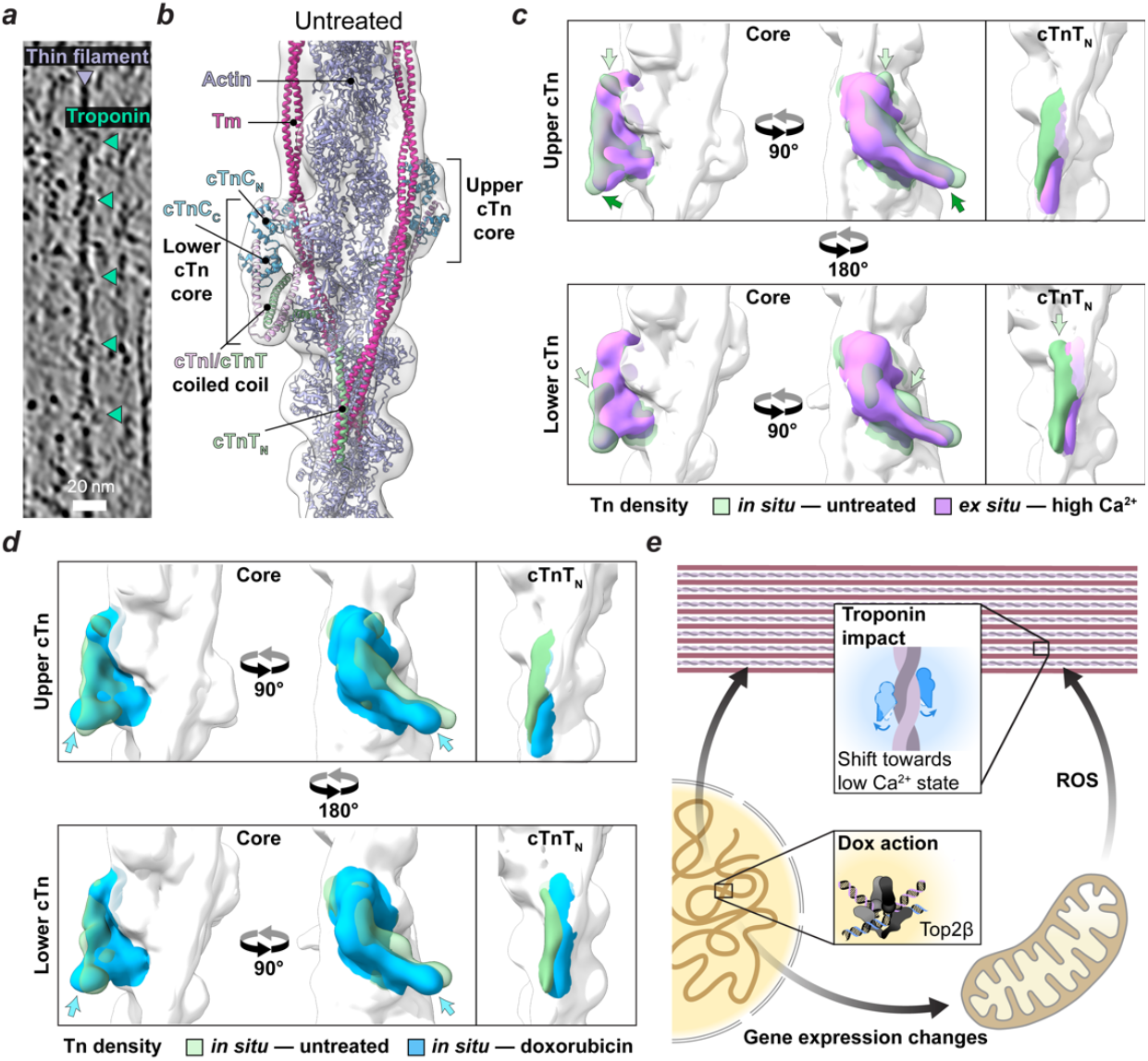
*In situ* structures of the troponin complexes are impacted by cardiotoxic drug treatment. ***(a)*** A tomogram slice including a thin filament (purple) with the troponin complexes bound (green). ***(b)*** The structure of the thin filament, reconstructed from untreated control cells, with the troponin complexes. A model of the troponin complex (PDB:6KN8) was docked into both sites, including cTnC (blue), cTnI (pink), and cTnT (green). ***(c)*** Several aspects of the *in situ* reconstruction from untreated control cells differ from ex situ. The untreated in situ maps of troponin (transparent green) are overlaid with the *ex situ* map from high calcium conditions (purple). The maps of the core complex (left) and cTnT N-terminal helix (right) are shown for the upper complex (top) and lower complex (bottom). ***(d)*** Sub-lethal doxorubicin exposure alters the troponin complex structures. The maps of untreated control cells (transparent green) and doxorubicin-treated cells (light blue) are overlaid, using the same format as in (c). ***(e)*** Our optimized cryo-ET platform reveals indirect effects of sub-lethal doxorubicin exposure on troponin structure. The primary target of doxorubicin (Dox) is thought to be Topoisomerase-IIβ (Top2β) in the nucleus. Altered gene expression patterns impact myofibril protein components and contribute to an increase in reactive oxygen species production by the mitochondria, which may also directly impact myofibril protein structure. We observe a shift towards the low calcium state of the troponin complex as a consequence of these global changes.

The troponin complex is composed of the Ca^2+^-binding subunit troponin C (cTnC), the regulatory subunit troponin I (cTnI), and the tropomyosin binding subunit troponin T (cTnT) (Fig. 4b). The troponin complex is known to assemble at two adjacent sites on the thin filament (the upper and lower complexes) (Fig. 4b). In response to Ca^2+^ binding at cTnC, the troponin complex shifts the position of tropomyosin to uncover myosin binding sites and initiate contraction^55^. Recent studies using reconstituted or isolated thin filaments have shown that the upper and lower troponin complexes can adopt a combination of low and high Ca^2+^ states at various Ca^2+^ concentrations^54,56^. However, the *in situ* structure of the troponin complex in cardiac cells is unknown.

In the thin filament model that docks best into our reconstruction, Ca^2+^ is bound to both troponin complexes (EMD:0729)^54^ (Fig. 4b, Fig. S5a). While our *in situ* reconstruction is similar to *ex situ* reconstructions with both troponin complexes in a high Ca^2+^ state^54,56^, several differences exist (Fig. 4c, Movie S4). In the upper complex, the core has a noticeable clockwise rotation and upward pitch (Fig. 4c, upper left, dark green arrows) and a protrusion emerges near the switch helix of cTnI in the model (Fig. 4c, upper left, light green arrows). The protrusion could reflect decreased flexibility of cTnI residues C-terminal to the switch helix or an alternate position of the cTnI switch helix on the cTnC_N_ surface. Interestingly, a similar density protrusion was observed in isolated thin filaments^56^ (PDB: 7KON)(Fig. S6). In the lower complex, there is an apparent loss of density at cTnC and an increase in density above the coiled-coil of cTnI/cTnT (Fig. 4c, lower left, light green arrows). Accompanying this, the N-terminal helix of cTnT shifts on the actin filament surface (Fig. 4c, lower right, light green arrows). This shift goes beyond the described position of cTnT in the high Ca^2+^ state^54^, and may represent an alternative interaction that influences functional coupling between neighboring tropomyosin strands.

By reconstructing the troponin complex from its cellular environment, we gained new structural insights into the breadth of conformations adopted by this central regulator of contraction. This further motivates an *in situ* structural biology approach for studying cardiac biology. Given our ability to reconstruct the troponin complex *in situ* and at a high-enough resolution to track functional changes, we next determined the extent that troponin complex structures respond to perturbations, using the cardiotoxic drug doxorubicin and the *MYH7*^WT/G256E^ mutation associated with hypertrophic cardiomyopathy.

### Changes in troponin structure are part of the systemic effects of doxorubicin-induced cardiotoxicity

Doxorubicin is an anti-cancer drug with a known side-effect of cardiotoxicity. The primary target for doxorubicin-induced death in cardiomyocytes is thought to be Topoisomerase-Iiβ (Top2β) in the nucleus^57^.

However, subcellular damage from doxorubicin has been associated with multiple phenotypes, including sarcomeric disarray and deterioration of myofibrils^58–60^. The mechanisms driving myofibrillar deterioration and contractile dysfunction^61^ during doxorubicin treatment are not clearly understood and are likely indirect, possibly due to oxidative injury from pathological mitochondrial metabolism^59^. Given our incomplete understanding of the molecular pathways of doxorubicin-induced cardiotoxicity, and the apparent complexity of the phenomenon, we recognized that our platform provided a unique opportunity to determine the structural effects of doxorubicin on sarcomeric structure at high resolution. As a proof-of-principle, we determined the impact of sub-lethal doxorubicin concentrations on troponin complex structures in hiPSC-CMs.

We treated hiPSC-CMs with 100 nM doxorubicin for 24 hours before processing and imaging cells. Drug-treated cells were split from the unperturbed hiPSC-CMs immediately before treatment and samples were processed in parallel to control for variables like genetic background and maturity. The concentration and duration of treatment were chosen because they were known to be below the LD_50_ for hiPSC-CMs but sufficient to induce gene expression changes and alter contractility^58^. The thin filament with troponin was reconstructed from drug-treated cells to a global resolution of 12.7 Å (Fig. S2b).

Comparing troponin structures in doxorubicin-treated versus untreated hiPSC-CMs revealed several interesting changes (Fig. 4d). The distinct features of the untreated *in situ* troponin complexes are removed in doxorubicin-treated cells, including the density protrusion at the TnI switch helix in the upper complex, the density rearrangement in the lower troponin complex core, and the shift in the lower TnT helix (Fig. 4d). Doxorubicin treatment could alter the environment of troponin complexes and reduce the influence of local interactions that contribute to the unique features we observed in untreated cells. At the same time, both troponin complexes rotate clockwise relative to the untreated control (Fig. 4d blue arrows, Fig. S5e, Movie S5). This rotation in doxorubicin-treated cells is in the same direction as the low Ca^2+^ *ex situ* reconstructions, but at a smaller magnitude (Fig. S5b). These conformational shifts are consistent with the lowered contractility^57,58,61^ phenotype of doxorubicin treatment.

Doxorubicin-dependent changes in the troponin complex demonstrate the potential of our *in situ* approach for revealing structural changes that depend on complex and indirect cellular processes (Fig. 4e). Expanding our approach to other sarcomeric proteins such as myosin will be helpful in understanding the detailed changes leading to contractile dysfunction, while reconstructing key macromolecules outside of the sarcomere will be important for dissecting the broader mechanisms by which doxorubicin exerts its systemic effects.

### Systemic effects of a hypertrophic cardiomyopathy-associated mutation include reduced occupancy of the troponin complex

Inherited cardiomyopathies are a major cause of heart failure^62^, and a better understanding of the mechanisms responsible could improve preventative diagnostics and lead to new treatments. While the mutations leading to hypertrophic cardiomyopathy can occur in multiple genes encoding sarcomere components, mutations in the thick filament make up the plurality of cases^63^. In recent work, an hiPSC-CM cell line with a mutation in the heavy chain of myosin (β-MHC G256E)^46^ was generated and characterized for disease phenotypes. The *MYH7*^WT/G256E^ genetic background was demonstrated to have the hypercontractility, altered gene expression, and increased metabolism that are considered hallmarks of hypertrophy at the cellular level^46^. However, the broader effects of the mutation on sarcomere structure remain undetermined, as is the case for many inherited cardiomyopathies. To test the potential of an *in situ* structural approach to answer these questions, we determined the impact of a *MYH7* ^WT/G256E^ background on troponin structure.

We started with the hiPSC cell line *MYH7*^WT/G256E^ (mutant) and its isogenic wild-type background, *MYH7*^WT/WT^ (isogenic control). Both cell lines carry an EGFP fluorescent tag at the endogenous locus of α-actinin, allowing for fluorescent visualization of sarcomeres (Fig. S1f). Cells were differentiated into CMs, processed, and imaged, using the same workflow as above but with two notable differences (Methods). First, the differentiated hiPSC-CMs matured for roughly 3 weeks longer in this experiment. Second, we used correlative light and scanning electron microscopy to select cells with well-organized myofibrils for lamellae, as evident in fluorescence images (Fig. S1f). Because of these differences, we expected the sarcomeres to be in a more mature state. The thin filaments with troponin were reconstructed at a global resolution of 12.4 Å and 18.9 Å for mutant and isogenic control, respectively (Fig. S2b).

As observed in the untreated hiPSC-CMs, the *in situ* troponin complex reconstructed from isogenic control cells was best fit by a model with both troponin complexes in a high Ca^2+^ state (Fig. 5a, Fig. S5c). However, we noted striking differences in the *in situ* architecture as compared to the *ex situ* model. In both the upper and lower complexes, there is a decrease in density at the N-terminal TnC domain and an increase in density at two locations— first, the interface between the C-terminal TnC domain and the TnI/TnT coiled-coil (Fig. 5a, fuschia arrows, Movie S6), and second, the tip of the TnI/TnT coiled coil (Fig. 5a, hot pink arrows, Movie S6). These additional densities could be explained by cross-bridging interactions with myosin, myosin binding protein C, or other components of the mature sarcomere^27^.

**Figure 5:**
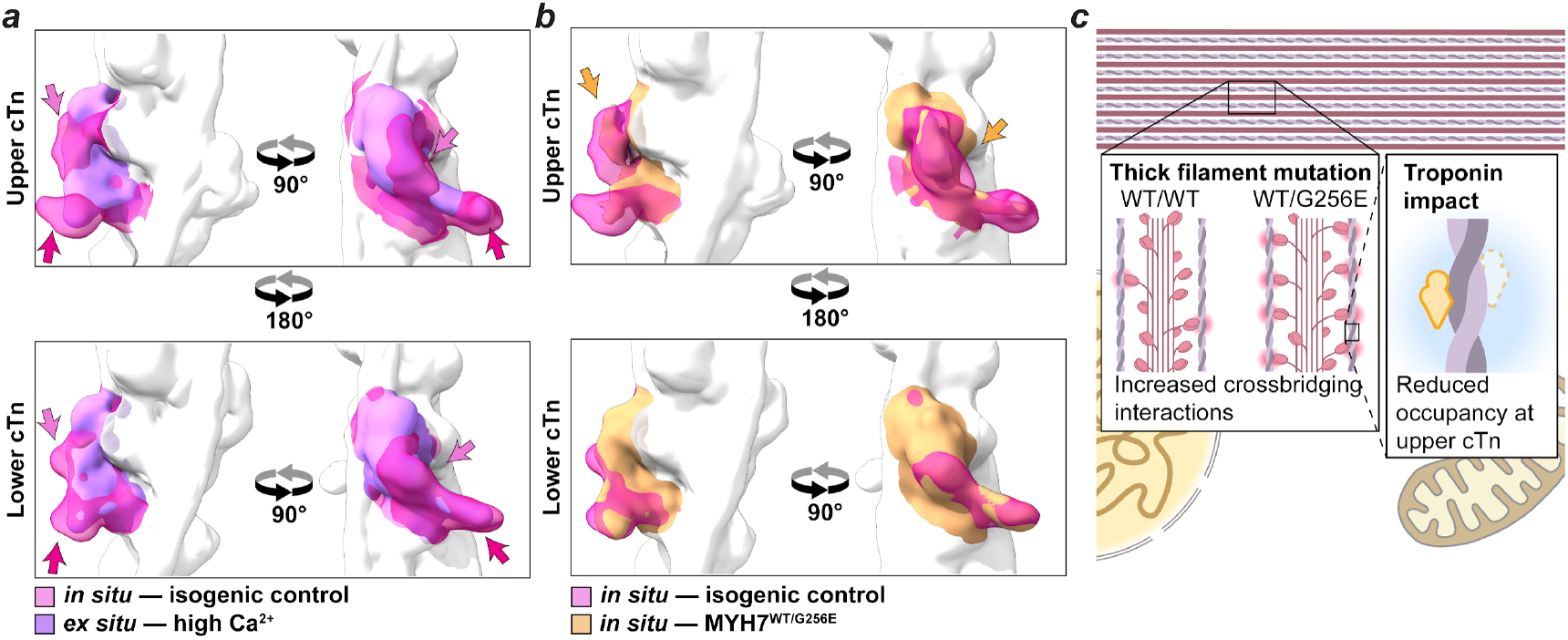
A mutation associated with hypertrophic cardiomyopathy alters troponin complex occupancy. ***(a)*** The *in situ* reconstruction from isogenic control cells differ from *ex situ*. The isogenic *in situ* maps of troponin (transparent hot pink) are overlaid with the *ex situ* map from high calcium conditions (purple). The maps of the core complex are shown for the upper complex (top) and lower complex (bottom). ***(b)*** The *MYH7* ^WT/G256E^ mutant background alters the troponin complex structures. The maps of isogenic control cells (transparent hot pink) and mutant cells (gold) are overlaid, using the same format as in (a). ***(c)*** The *MYH7* ^WT/G256E^ mutation has a hypercontractile phenotype associated with increased cross-bridging interactions in the myofilament. We observe reduced occupancy at the upper troponin complex in the mutant background.

Comparing troponin complex structures between *MYH7*^WT/G256E^ and its isogenic control, we were struck by a prominent loss of density in the upper complex of *MYH7*^WT/G256E^ (Fig. 5b, gold arrows, Movie S7). This density loss is consistent with partial occupancy of the upper troponin complex in the mutant background. Partial occupancy has been observed previously in *ex situ* studies. In myofibrils isolated from cardiomyocytes^56^, a sizable fraction of thin filament segments were missing either the upper or lower complex. Proteolysis of troponin complex subunits has also been observed under stress conditions^64–67^. Depletion of troponin complexes from the thin filament increases myofilament contractility^66,68,69^, which suggests that reduced occupancy *in situ* contributes to the cellular phenotypes associated with hypertrophic cardiomyopathy^46^. The induction of *TNNI3* gene expression in the *MYH7*^WT/G256E^ genetic background^46^ could represent a compensatory mechanism to counteract this exacerbating effect.

These results demonstrate the potential of in-cell structural biology to interrogate the systemic effects of gene variants associated with inherited cardiomyopathies. Our discovery of a structural state of troponin predicted to exacerbate the *MYH7* mutant phenotype illustrates the importance of an expanded structural characterization of the cellular system to fully understand mechanisms leading to disease states. It will be interesting to determine the extent to which the loss of troponin is generally associated with hypertrophic cardiomyopathy.

## Discussion

The regulatory pathways controlling our heartbeat are complex, spanning multiple spatial scales. This has made it difficult to reconstitute the full system using traditional biochemical approaches and hampered our full mechanistic understanding of cardiac pathologies. Here we describe the use of a customized cryo-ET platform applied to human induced pluripotent stem cell–derived cardiomyocytes (hiPSC-CMs) to tackle this challenge by directly determining structures of select cardiac proteins in cells down to sub-nanometer resolution. Our optimized platform incorporates multiple advances including micropatterned electron microscopy grids^37,38^ to control hiPSC-CM attachment, correlated cryogenic light and focused ion beam microscopy to prepare representative hiPSC-CMs lamellae for cryo-ET, and recently developed advances in data collection and image processing methods^39–45^. Using our platform, we have reconstructed 8 *in situ* structures of the human cardiac thin filament and measured the response of thin filament structure to both a cardiotoxic drug (doxorubicin) and a hypertrophic cardiomyopathy-associated mutation (*MYH7*^WT/G256E^).

Reconstructing the troponin complex from within cells revealed several notable differences from the canonical structural models. Some of these *in situ* differences have analogues from previous *ex situ* work, including a shift near the cTnI switch helix, reduced occupancy of the troponin complex, and partial rotation of the cTnI/cTnT coiled-coil^56^. Other changes have not been captured previously, like the density changes in the core complex and the shift of the cTnT N-terminal helix. The *in situ* reconstructions may be highlighting states that are of low population *ex situ* or that depend on a larger cellular context, like cross-bridging interactions, post-translational modifications, alternate isoforms, and proteolysis. By demonstrating the existence of these states as predominant members of the molecular population within cells, this work validates the physiological relevance of previous observations, introduces new conformational states not observed in *ex situ* experiments, and motivates future efforts to incorporate them into the models of troponin structure and function. The next step in characterizing these states will be to expand *in situ* data collection efforts. Increasing the number of thin filament segments analyzed will enable further classification and reconstruction of additional states. This could be particularly important for better quantifying effects like partial troponin occupancy in the *MYH7*^WT/G256E^ background. Increasing the resolution of the troponin complex reconstructions will also help us better define the molecular interactions behind these observed differences.

A major focus of this work was using our optimized platform to track the systemic effects of disease-associated perturbations on thin filament structure. Testing both doxorubicin exposure and *MYH7*^WT/G256E^, we observed changes in troponin reconstructions that were consistent with known cellular phenotypes of these perturbations. This is an important demonstration of the power of *in situ* structural biology to study cardiovascular disease, especially when the effects are indirect and context-dependent. In this study, we used a doxorubicin concentration and exposure time that had only mild effects on phenotypes like gene expression, metabolism, and myofibrillar organization^58^. Expanding this effort to a range of doxorubicin concentrations and exposure times will provide complementary structural information on disease progression. Structural changes may not only provide a diagnostic tool for changes in cellular physiology, but also generate new leads for treatment. We are particularly intrigued by the loss of troponin occupancy observed in the *MYH7*^WT/G256E^ genetic background. Reduced occupancy may arise from aberrant cross-bridging interactions that destabilize the complex or an induction of proteolysis^64–67^.

Loss of troponin is likely to exacerbate the hypercontractile phenotype of *MYH7*^WT/G256E^ cells^66,68,69^, which suggests that therapeutic interventions that prevent this loss could serve as an independent strategy for counteracting the disease phenotype. To determine the value of this strategy, it will be important to test the generality of the connection between troponin loss and hypercontractility across the breadth of genetic backgrounds associated with hypertrophic cardiomyopathy.

Fully realizing the potential of our approach will require new strategies to better integrate information at the molecular and cellular scales—from multiple molecules and across multiple compartments. We are particularly interested in sites beyond the sarcomere that are intimately involved in the progression of cardiovascular diseases, like protein complexes in the mitochondria that generate energy and Ca^2+^ handling proteins that control the contraction cycle. Incorporating more sophisticated model systems like engineered cardiac tissues into our platform will also be important for capturing structures from a more tissue-like state. The results from our work demonstrate the utility of our approach for dissecting previously unobserved structural states in the native context of cardiomyocytes and in response to cellular perturbations, laying the foundation for an exciting new approach to structural cardiac biology.

## Acknowledgements

We thank Gong-her Wu, Patrick Mitchell, Grigore Pintilie, Muyuan Chen, Andrew Wu, and members of the Chiu and Wu lab for helpful discussions. This research was supported by: T32 NIBIB 2T32EB009035 and NHLBI K99HL161392 (to R.A.W.), and other support including NIH S10OD021600, DOE (BERFWP 100463), Silicon Valley Community Foundation Chan Zuckerberg Initiative (2021-234593) (to W.C.). Some of this work was performed at the Stanford-SLAC Cryo-EM Center and the Stanford-SLAC Cryo-ET Specimen Preparation Center under the support of the National Institutes of Health Common Fund’s Transformative High Resolution Cryo-electron Microscopy Program (U24GM139166 and U24GM129541). The authors would also like to thank the following SLAC cryoEM personnel for their invaluable support and assistance: Patrick Mitchell, Megan Mayer, Corey Hecksel, Chensong Zhang and Lydia-Marie Joubert. Some of the work was performed at nano@Stanford labs, which are supported by the National Science Foundation as part of the National Nanotechnology Coordinated Infrastructure under award ECCS-2026822. M.N. was supported by NHLBI K99HL166773. This paper was typeset with the bioRxiv word template by @Chrelli: www.github.com/chrelli/bioRxiv-word-template.

## Author contributions

R.A.W. and W.C. conceived the project and designed the experiments. W.C. and R.A.W. supervised the project and acquired funding. M.N., A.S.V., and P.G. prepared hiPSC-CM samples under the supervision of J.C.W., J.A.S., and D.B. L.E. patterned electron microscopy grids under the supervision of A.R.D. with input from R.A.W., A.S.V., and M.N. R.A.W. processed samples and acquired data. R.A.W. processed and analyzed the data with help from G.C.M. in annotation and M.F.S. with initial interpretation. R.A.W. and W.C. wrote the manuscript draft with the help from all authors.

## Online Methods

### Micropatterning of EM grids

Digital files of rectangles with a 1:7 aspect ratio and areas of 2000 μm^2^ (16.903 μm by 118.32 μm) and 1500 μm^2^ (14.64 μm by 102.465 μm) were generated in *ImageJ*^70^ (Digital Files S1 & S2, pixel conversion is 3.58 pixels/μm).

EM grids (UltrAuFoil R2/2 Au 200 mesh Q250AR2A and Quantifoil R2/2 Au 200 mesh Q2100AR2) were micropatterned using the Alvéole PRIMO micropatterning system^38,71,72^. Unpatterned grids were placed holey film–up on silicone sheeting (Specialty Manufacturing Inc.) on glass coverslips to prevent them from moving during exposure to plasma and incubation. They were exposed to atmospheric plasma at 22 Watts for 10 seconds using a benchtop plasma etching system (Plasma Etch PE50). The glass coverslip containing the grids was placed on Parafilm (Bemis™ PM999) and several milliliters of 0.01% Poly-L-Lysine (CAS Number 25988-63-0, p4707 Sigma) were added to the surface immediately following the plasma exposure. After 1 hour at room temperature, the grids were rinsed three times in deionized water (DIW) and incubated in a 100 mg/mL solution of mPEG-Succinimidyl Valerate, MW 5,000 (PEG-SVA, Laysan Bio Inc) in 0.1 M HEPES (pH 8.5) for 1 hour. They were then rinsed several times in DIW.

Each grid was lifted from the silicone, blotted dry, and held in negative pressure tweezers while 3 μL of a 1:6 solution of photoinitiator (PLPP gel, nanoscaleLABS, LLC) in pure ethanol was added to the surface. Following drying of the PLPP gel, the grids were placed holey film–down on a glass coverslip and exposed to UV in the Alvéole PRIMO micropatterning system at 50 mJ/mm^2^. They were then rinsed in DIW several times and stored in 15 μL droplets of DIW contained to custom silicone wells in glass bottom dishes until protein backfill.

### hiPSC-cardiomyocyte differentiation and re-plating to EM grids

#### *MYH7*^WT/WT^ and *MYH7*^WT/G256E^ hiPSC-CMs

The WTC hiPSC cell line was generated by the Bruce R. Conklin Laboratory at the Gladstone Institutes and University of California– San Francisco (UCSF)^73^. Cells were maintained as hiPSCs and differentiated into cardiomyocytes, using methods described previously^36,74^ and available from the Allen Institute for Cell Science webpage (http://allencell.org). AICS-0097-113 (*ACTN2-mEGFP MYH7*^WT/WT^) and AICS-0097-141 (*ACTN2-mEGFP MYH7*^WT/G256E^) were developed via CRISPR/Cas9 gene editing and subjected to quality control at the Allen Institute for Cell Science^46^.

hiPSCs were differentiated into cardiomyocytes using an established small molecule protocol^75^ with modifications^36^. Briefly, hiPSCs were seeded onto Matrigel-coated 6-well tissue culture– treated plates and maintained for 4 days with mTeSR1 media (85850, Stemcell Technologies). On day 0, hiPSCs were induced with 7.5 μM CHIR99021 (13122, Cayman Chemical) in RPMI 1640 (11875-093, Gibco) and 1X B27 minus insulin (A1895601, Gibco). After 48 hours, the media was replaced with 7.5 μM IWP2 (3533, Tocris) in RPMI 1640 and 1X B27 minus insulin. On day 4, the media was replaced with RPMI 1640 and 1X B27 minus insulin. From day 6 onwards, differentiated cells were maintained in RPMI 1640 and 1X B27 plus insulin (A1895601, Gibco) with media changes every 2-3 days. RPMI 1640 contains 420 μM Ca^2+^, which is sufficient to support the contraction cycle. After the onset of beating (∼day 10) cells were grown in RPMI media without glucose with B27 for 4 days and cells were replated on day 15 for subsequent growth and maturation.

On day 48, cells were plated onto the micropatterned electron microscopy grids by lifting with TrypLE and resuspending in replating media (RPMI + B27 with 10% Knock-out Serum Replacement and 10 μM Rock inhibitor) at a density of 100,000 cells/mL. This was done for both AICS-0097-113 and AICS-0097-141 hiPSC-CMs expressing an endogenous fluorescent tag of the sarcomere protein α-actinin. The micropatterned grids were held in place in a glass bottom dish with a custom PDMS sticker which enabled the use of small volumes (10-50 μL) of solutions for incubation. After patterning, the grids were sterilized by a 10 min incubation in 70% ethanol, subsequently rinsed in PBS -/-and incubated with a 1:150 dilution of Matrigel in DMEM/F12 media for 1 hour. After removal of the Matrigel solution, 10 μL of replating media were added to each grid, and then approximately 500 cells were added to each grid in 5 μL of resuspension. After 2 hours of attachment, an additional 2 mL of replating media were added to each plate, and then media was replaced with RPMI + B27 after 2 days and cells were cultured for 7-10 days to recover before vitrification. We visually verified that the hiPSC-CMs were beating on the micropatterned EM grids during recovery. Cells were not synchronized before vitrification.

#### Doxorubicin treated hiPSC-CMs

Human iPSCs were generated from peripheral blood mononuclear cells collected from healthy donors using Sendai virus and cryopreserved at Stanford CVI iPSC Biobank (SCVI-273). Human iPSCs were maintained in Essential 8 (E8) medium (Thermo Fisher Scientific, A1517001) on Matrigel-coated (1:200 dilution, Corning, 356231) 6-well plates. For passaging of hiPSCs, hiPSCs were detached with 0.5 mM EDTA and cultured in E8 medium with 10 μM Y-27632 ROCK inhibitor (Selleck Chemicals, S1049) for 24 h. Cardiomyocyte differentiation was carried out in RPMI 1640 (Thermo Fisher Scientific, 11875-119) with B27 supplement minus insulin (Thermo Fisher Scientific, A1895601) when hiPSCs (at passages 20 - 35) reached 80 - 90% confluency, as previously described^36,58,75^. Briefly, hiPSCs were treated with 8 μM CHIR-99021 (Selleck Chemicals, S2924) for 2 days, followed by 1 day recovery, and then 5 μM IWR-1 (Selleck Chemicals, S7086) for another 2 days. Starting from day 7, the differentiation medium was changed to RPMI 1640 with insulin B-27 Supplement (Thermo Fisher Scientific, 17504044). From day 12, non-cardiomyocytes were removed by glucose starvation using RPMI without glucose (Thermo Fisher Scientific, 11879-020) with B27 supplement for 4 days. Patterned EM grids were prepared for cell attachment using the protocol as above. The purified cardiomyocytes on day 20-30 were replated using TripLE Select (Thermo Fisher Scientific, A1217701) to micropatterned cryo-EM grids at a density of 750 cells per grid. The hiPSC-CMs were cultured on the micropatterned grids for 5 days in RPMI 1640 containing 420 μM Ca^2+^, which is sufficient to support contraction. We visually verified that the hiPSC-CMs were beating on the micropatterned EM grids during the recovery period. During the last 24 h before vitrification, a subset of grids were treated with 100 nM doxorubicin. This treatment regimen was chosen as non-lethal while still impacting hiPSC-CM physiology and gene expression^58^. Cells were not synchronized before vitrification.

### Vitrification of cardiomyocytes, cryo-CLEM, and cryoFIB-SEM milling

Cells were vitrified by plunge freezing into liquid ethane using a Leica EM GP2 plunger. The grids were blotted from the opposite side of the holey film layer at 25 °C and 95% humidity.

Cells expressing an endogenous fluorescent tag on α-actinin were imaged using the integrated fluorescent light microscope (iFLM) in a cryoFIB-SEM (ThermoFisher Aquilos2 cryoFIB-SEM) to identify cells with ordered sarcomeric arrangement. In early experiments, before the integration of a fluorescent microscope into the cryoFIB-SEM chamber of the Aquilos, some grids were visualized using a Zeiss Airyscan2 (LSM800) confocal microscope with Linkam cryostage.

The vitrified cells were then milled to a thickness of ∼150 nm using a cryo-FIB, either manually or using the AutoTEM Cryo software (ThermoFisher). The milling was done using a ThermoFisher Aquilos1 or Aquilos2 cryoFIB-SEM with iFLM, following previously described workflows^76,77^. In brief, the grids were sputter coated with metallic platinum using an in-chamber plasma coater to minimize charging. The samples were then coated with a ∼500 nm organometallic platinum layer using a gas injection system and additionally sputter coated with a thin layer of platinum. SEM imaging was performed using 2-5 kV and 13 pA current to check milling progress. Milling was performed using a 30 kV ion beam while progressively decreasing the beam current from 300 pA to 30 pA. Micro-expansion joints were used during milling to avoid lamella bending and breakage^78^. Final polishing was performed manually. After polishing, some of the grids were coated with a thin layer of platinum.

### Cryo-ET data acquisition, image processing, and tomogram reconstruction

Cryo-ET datasets were acquired at 300 kV using Titan Krios transmission electron microscopes (ThermoFisher) equipped with K3 cameras and energy filters (Gatan) (Table S1). Data acquisition was performed using SerialEM^79^ software. Overview images of the lamellae were collected at 1.04 nm pixel size to identify areas for high-magnification tilt-series data collection. Tilt series projections were acquired at about 2.13 Å pixel size (42,000x nominal magnification) using a dose-symmetric^80^ data acquisition scheme with 2° or 3° increments starting from 0° or relative to the lamella pre-tilt. The images were recorded as movies divided into 10 frames. The total dose applied to the sample ranged from 105–120 e^-^/Å^2^. For datasets collected using K3 CDS mode, the targeted dose rate on the camera was 5–6 e^-^/pix/s; for non-CDS mode datasets, the targeted dose rate was 10–15 e^-^/pix/s.

Individual tilt frames were motion corrected using MotionCor2^40^. The tilt-series were then aligned and reconstructed with dose weighting using AreTomo^41^. The reconstructions were used to select tomograms without crystalline ice, with the correct field of view, and with sufficient contrast. This resulted in a final data set of 12 tomograms (isogenic control), 8 tomograms (mutant cells), 10 tomograms (untreated control), and 14 tomograms (drug treated cells). For better visualization, CTF-corrected and denoised tomograms were reconstructed using IsoNet^43^ with CTF estimates from Warp^39^. We trained a neural network model that can be used to denoise our full dataset using three representative tomograms. Denoised tomograms were only used for visualization and particle picking, and they were not used in the sub-volume averaging pipeline.

### Subvolume averaging

#### Particle identification

3D convolutional neural network (CNN)–based models were trained to recognize five classes of cellular features: thick filaments, thin filaments, membranes, ribosomes, and background using DeepFinder^42^ (Fig. S1b, Movie S1). Each macromolecule was manually annotated in 4-5 denoised tomograms down sampled by a factor of 4. We did not differentiate between myosin-bound and unbound segments of the thin filament in the manual annotations, and we see continuous labeling of the thin filament in our annotation, showing that the model does not exclude myosin-bound sections. Manually annotated coordinates were used to train two CNN models to recognize the different classes. One model was trained on data collected with 2° tilt increments, and the other was trained on data collected with 3° increments. The trained models were used to annotate and pick positions of the thin filament in all of the tomograms. In the isogenic control dataset, 92,224 segments of the thin filament were picked from 12 tomograms with an inter-segment distance of 51 Å. In the mutant dataset, 95,513 segments were picked from 8 tomograms with an inter-segment distance of 68 Å. In the untreated control dataset, 40,621 segments were picked from 10 tomograms with an inter-segment distance of 51 Å. Finally, in the drug-treated dataset, 30,249 segments were picked from 14 tomograms with an inter-segment distance of 85 Å. Particles with overlapping segments—defined as an inter-segment distance of less than 60 Å—were filtered from analysis during reconstruction, so we do not expect variations in the inter-segment distances used for particle picking in the different experimental conditions to significantly affect the results. Segments picked for reconstruction came from both the A-band and I-band regions of thin filaments.

#### Thin filament refinement and classification

Sub-volume averaging of the thin filament was performed in RELION 4.0^44^ and Warp^39,81^/M^45^. The motion-corrected tilts and tilt series alignment parameters from AreTomo were imported into Warp, where the tilt-series CTFs were estimated. Pseudo-subtomograms were extracted in RELION using Warp’s CTF estimates. Subsets of the data were used for *de novo* initial model generation with C1 symmetry for each of the datasets. The full datasets were then aligned to and refined using the reference and subjected to 3D classification. Selected particles (16,451, 15,147, 9,851, and 7,555 for isogenic control, mutant, untreated control, and drug treated, respectively) were further refined in RELION and the results were imported into M. In M, the image-space and volume-space warping grids, the particle poses, and the stage angles were refined in 6 iterations. Starting with iteration 4, CTF parameters were also refined. The final global resolutions were 8.7 Å, 9.3 Å, 9.1 Å and 9.5 Å, respectively, for isogenic control, mutant, untreated control, and drug treated.

To reconstruct the thin filament with the troponin complex, the subvolumes were recentered towards the cross-point between the 36 nm repeats of the thin filament. After refinement and classification in RELION, the following number of particles were selected for each dataset: 1,463 (isogenic control), 2,114 (mutant), 721 (untreated control), and 2,737 (drug treated). After refinement in M, the final global resolutions for these datasets were 19.5 Å (isogenic control), 12.4 Å (mutant), 18.0 Å (untreated control), and 12.4 Å (drug treated).

### Visualization and figure preparation

To better visualize the subcellular arrangement of the variety of macromolecules in cardiomyocytes (Fig. 1a, Fig. 2c), the macromolecules were semi-automatically annotated using the convolutional neural network algorithm in EMAN2^82^. The segmented surfaces were then visualized in UCSF ChimeraX^83^. Tomographic slices used to visualize the myofibrils were prepared in 3dmod^84^.

To compare the *in situ* reconstructions to each other and *ex situ* maps, the maps were low-pass filtered to a common resolution close to the lowest resolution map within the set. For the thin filament without troponin this was 10 Å, for the thin filament with troponin, this was 18 Å. Low-pass filtering was performed using EMAN2^82^. All figures with subvolume-averaged maps were created using UCSF ChimeraX^83^. ChimeraX and Napari^85^ were used to visualize the tomograms and prepare movies. We used the *dock_in_map* function of PHENIX^86^ to do a rigid fit of *ex situ* models into each of the maps.

## Data availability

Cryo-ET density maps generated in this study will be available in the EM Data Bank with the following accession codes: EMD-43944 (*in situ* structure of the cardiac thin filament without troponin from isogenic control (*MYH7*^WT/WT^) hiPSC-CMs), EMD-49349 (*in situ* structure of the cardiac thin filament with troponin from isogenic control (*MYH7*^WT/WT^) hiPSC-CMs), EMD-49346 (*in situ* structure of the cardiac thin filament without troponin from mutant (*MYH7*^WT/G256E^) hiPSC-CMs), EMD-49351 (*in situ* structure of the cardiac thin filament with troponin from mutant (*MYH7*^WT/G256E^) hiPSC-CMs), EMD-49345 (*in situ* structure of the cardiac thin filament without troponin from untreated hiPSC-CMs), EMD-49350 (*in situ* structure of the cardiac thin filament with troponin from untreated hiPSC-CMs), EMD-49348 (*in situ* structure of the cardiac thin filament without troponin from doxorubicin-treated hiPSC-CMs), EMD-49352 (*in situ* structure of the cardiac thin filament with troponin from doxorubicin-treated hiPSC-CMs). Publicly available entries used in this study are EMD-0728/PDB 6KN7 (B-state), EMD-0729/PDB 6KN8 (C-state), EMD-13996 (M-state), and PDB 7KON (shift of cTnI switch helix in Fig. S6).

## Ethics statement

Cell lines AICS-0097-113 *ACTN2-mEGFP MYH7*^WT/WT^ and AICS-0097-141 *ACTN2-mEGFP MYH7*^WT/G256E^ were derived from WTC hiPSC by the Allen Institute for Cell Science. The WTC hiPSC line was obtained from a healthy adult by the Bruce R. Conklin Laboratory at Gladstone Institute and UCSF. Informed consent was obtained from the subject. The hiPSC cell line used for doxorubicin treatment was obtained from the Stanford CVI BioBank. Stem cells were derived from patient blood samples, and consent for research was obtained for all patients under Stanford IRB 29904.

## Supplementary Figures

**Figure S1:**
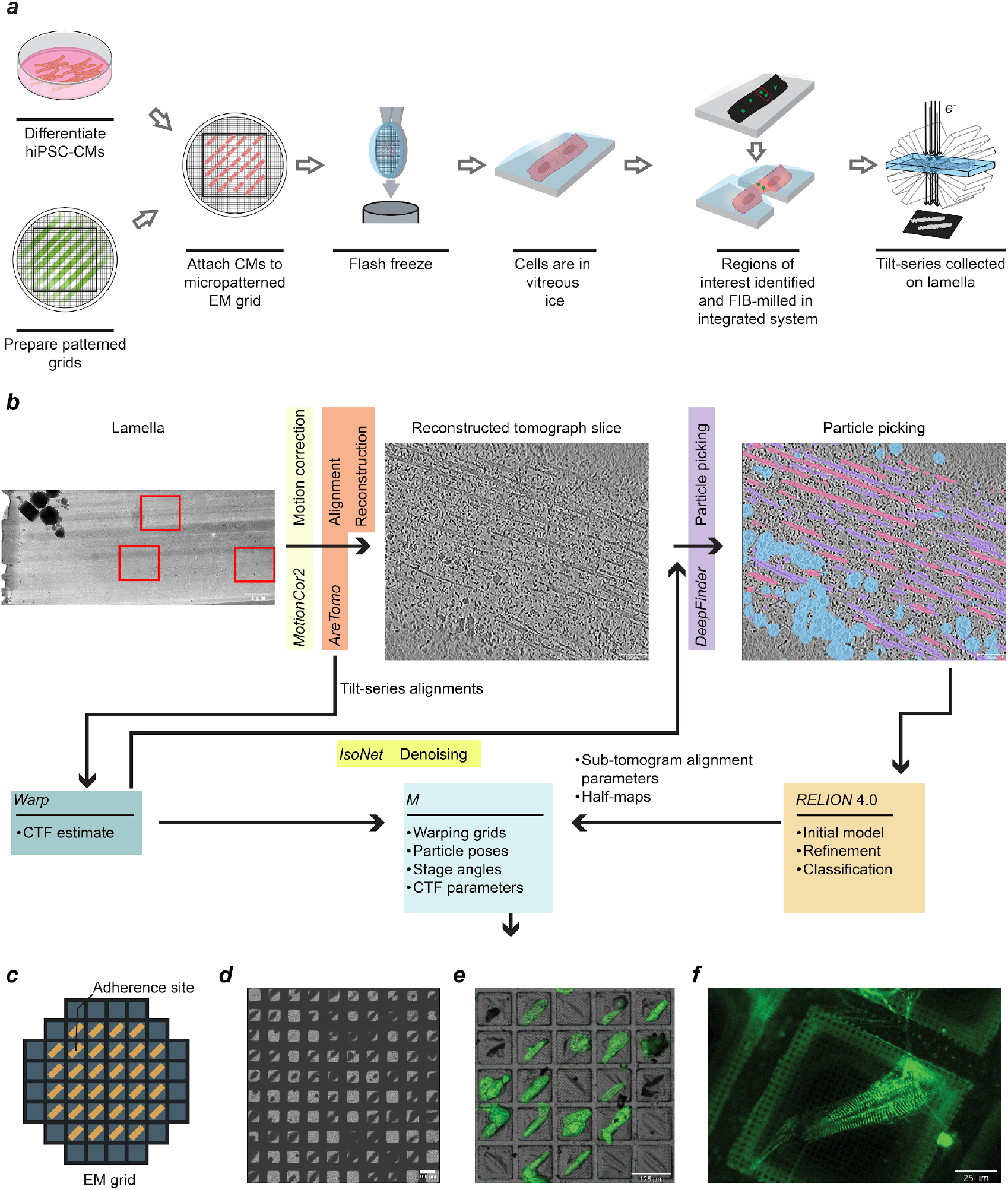
An optimized experimental and computational platform for visualizing hiPSC-cardiomyocytes using cryo-ET. **(*a*)** Human induced pluripotent stem cells (hiPSCs) are differentiated into cardiomyocytes (hiPSC-CMs) then attached to micropatterned EM grids. Once attached to the grid, cells are flash frozen to encase in vitreous ice. All subsequent steps are performed at cryogenic temperatures. Regions of interest within the cells are identified using correlated light and electron microscopy, then milled using a focused ion beam to create thin lamellae. Finally, a tilt-series is collected in the transmission electron microscope for subsequent reconstruction as a 3-dimensional tomogram. **(*b*)** A schematic of the data processing workflow, as detailed in the **Methods**. Tilt series were collected on regions of interest (red boxes within the lamella), MotionCor2 was used for motion correction, and AreTomo was used for alignment and reconstruction of the 3D tomogram. Alignments from AreTomo were passed to Warp to generate a CTF estimate that was passed to IsoNet to denoise the tomogram. Particle picking was performed in denoised tomograms using DeepFinder, and particle coordinates were passed to RELION 4.0. In RELION, an initial model was generated then refined, followed by particle classification. Sub-tomogram alignment parameters and half-maps were then passed to M for further refinement. **(*c*)** The diagonal rectangle pattern (ratio 7:1) used to guide hiPSC-CM adherence and shape. **(*d*)** A cryo-TEM atlas showing cells attached to a micropatterned EM grid. **(*e*)** Light/fluorescent microscopy overview of hiPSC-CMs expressing fluorescently tagged α-actinin. Cells are attached to a micropatterned grid to guide hiPSC-CM attachment and shape. **(*f*)** High-magnification fluorescent imaging within the FIB-SEM is used to identify sarcomeres and target regions for lamella generation.

**Figure S2:**
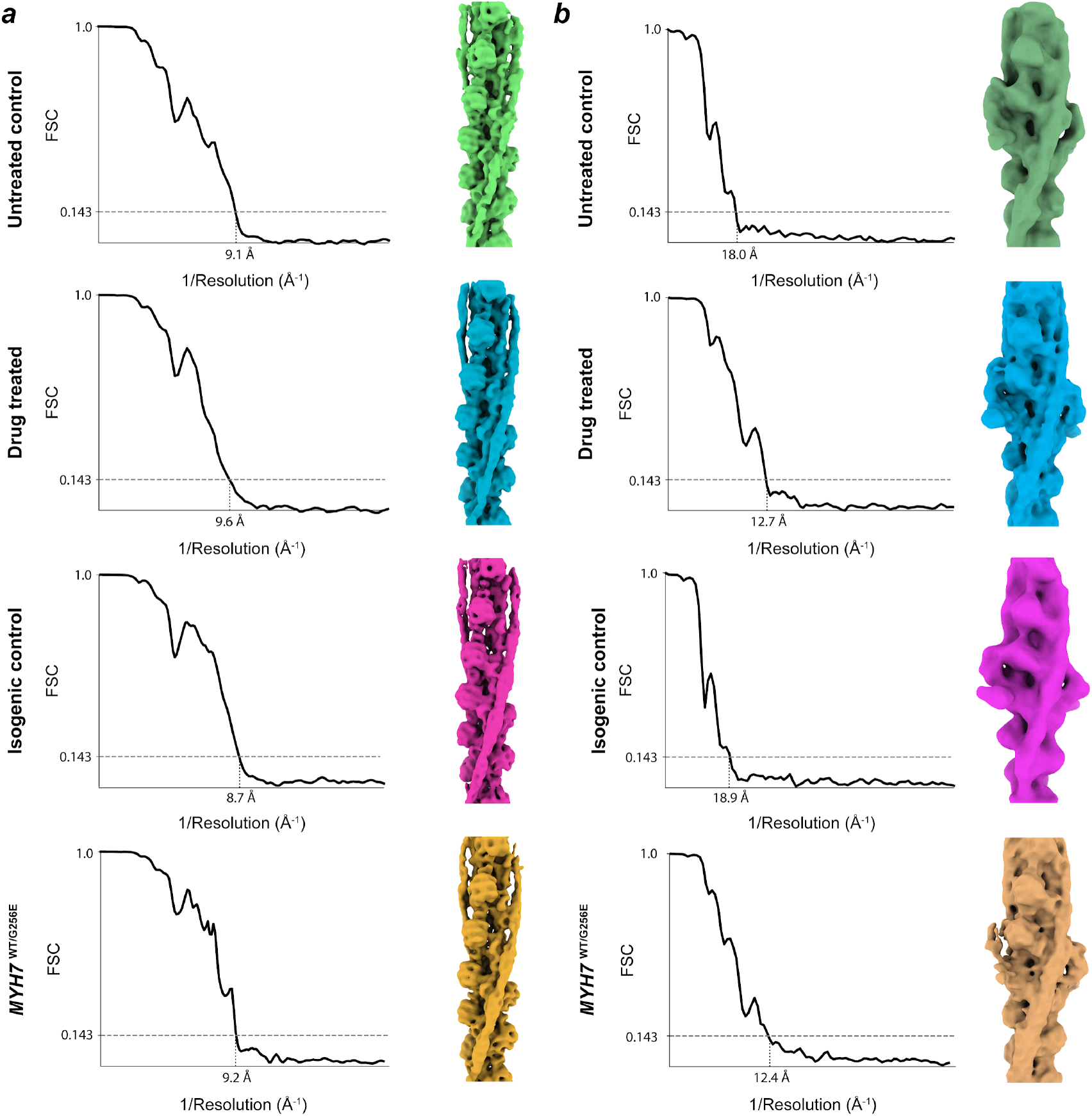
Fourier shell correlation plots of the reconstructed maps. **(*a*)** Fourier shell correlation (FSC) plots are shown for the thin filaments reconstructed from untreated control (top), drug-treated (doxorubicin, second from top), isogenic control (second from bottom), and *MYH7*^WT/G256E^ (bottom). The corresponding maps are shown to the right. **(*b*)** Fourier shell correlation (FSC) plots are shown for the thin filaments, with troponin included, using the same layout as in ***a***

**Figure S3:**
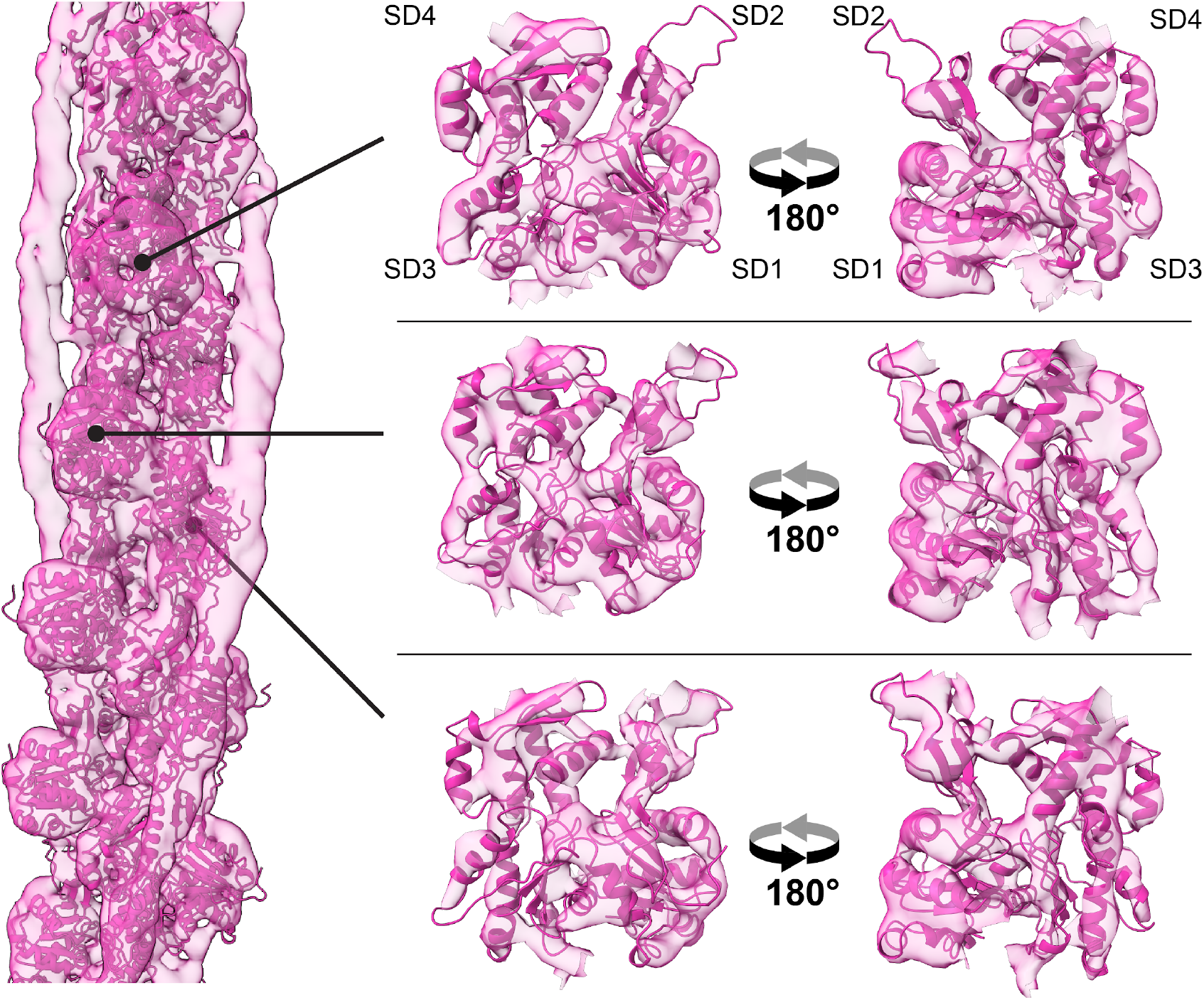
*In situ* structure of the thin filament resolves actin helices at 8.7 Å global resolution. (Left) An *in situ* structure of the thin filament reconstructed from the isogenic control cells (Right) 3 Individual actin monomers viewed from the front and back, with the 4 subdomains (SD1-4) labeled for the top monomer. The map resolves individual helices of the actin monomers.

**Figure S4:**
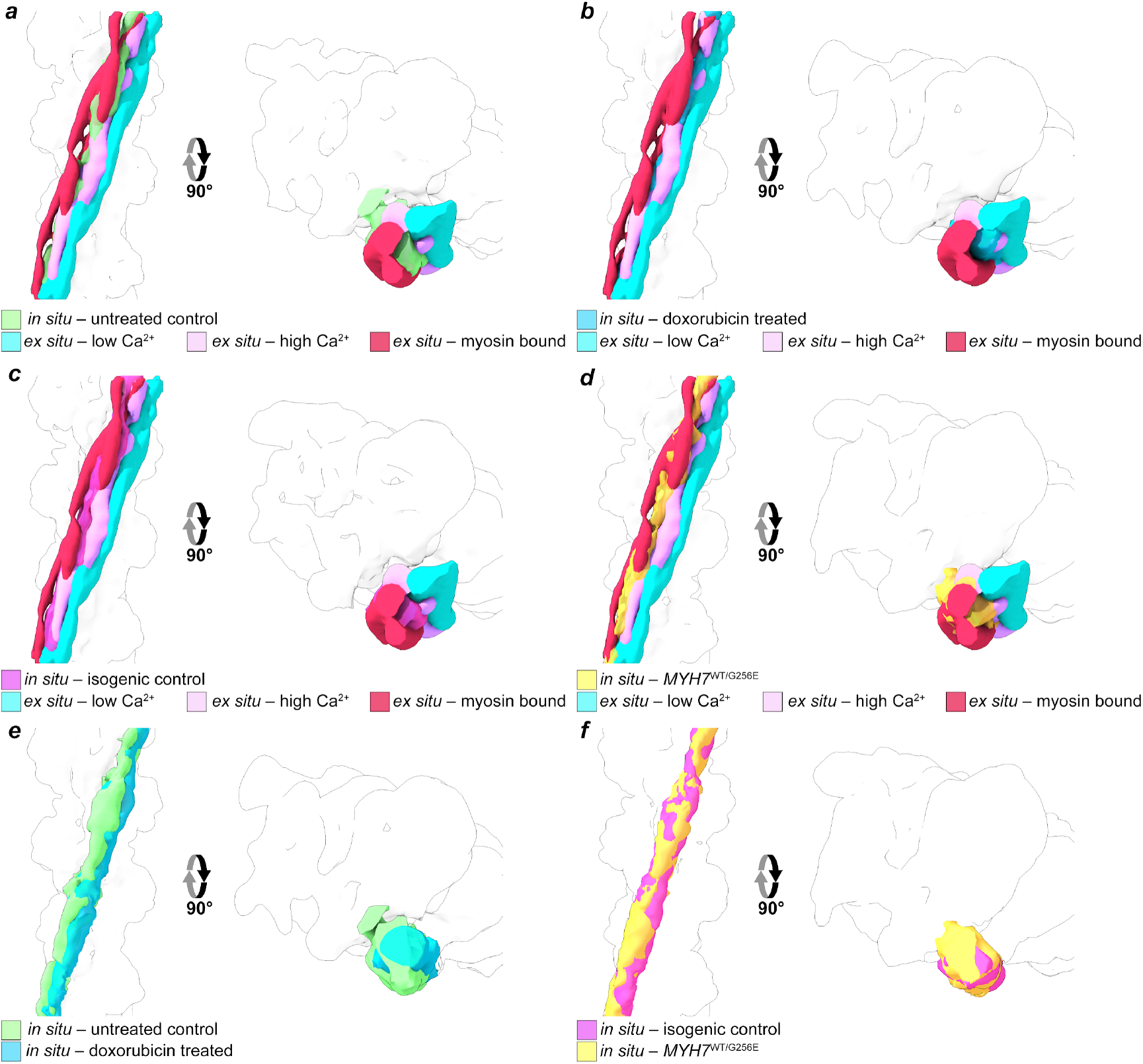
*In situ* positions of tropomyosin. Two views of the reconstructions of tropomyosin from different sources. **(*a*)** The *in situ* reconstruction from untreated control cells (green) compared to the *ex situ* low Ca^2+^ state (cyan, EMD-0728), the *ex situ* high Ca^2+^ state (purple, EMD-0729), and the *ex situ* myosin-bound state (red, EMD-13996). **(*b*)** The *in situ* reconstruction from doxorubicin-treated cells (blue) compared to the *ex situ* low Ca^2+^ state (cyan), the *ex situ* high Ca^2+^ state (purple), and the *ex situ* myosin-bound state (red). **(*c*)** The *in situ* reconstruction from isogenic control cells (hot pink) compared to the *ex situ* low Ca^2+^ state (cyan), the *ex situ* high Ca^2+^ state (purple), and the *ex situ* myosin-bound state (red). **(*d*)** The *in situ* reconstruction from *MYH7*^WT/G256E^ cells (gold) compared to the *ex situ* low Ca^2+^ state (cyan), the *ex situ* high Ca^2+^ state (purple), and the *ex situ* myosin-bound state (red). **(*e*)** The *in situ* reconstruction from untreated control cells (green) compared to the doxorubicin-treated state (blue). **(*f*)** The *in situ* reconstruction from isogenic control cells (hot pink) compared to the *MYH7*^WT/G256E^ state (gold).

**Figure S5:**
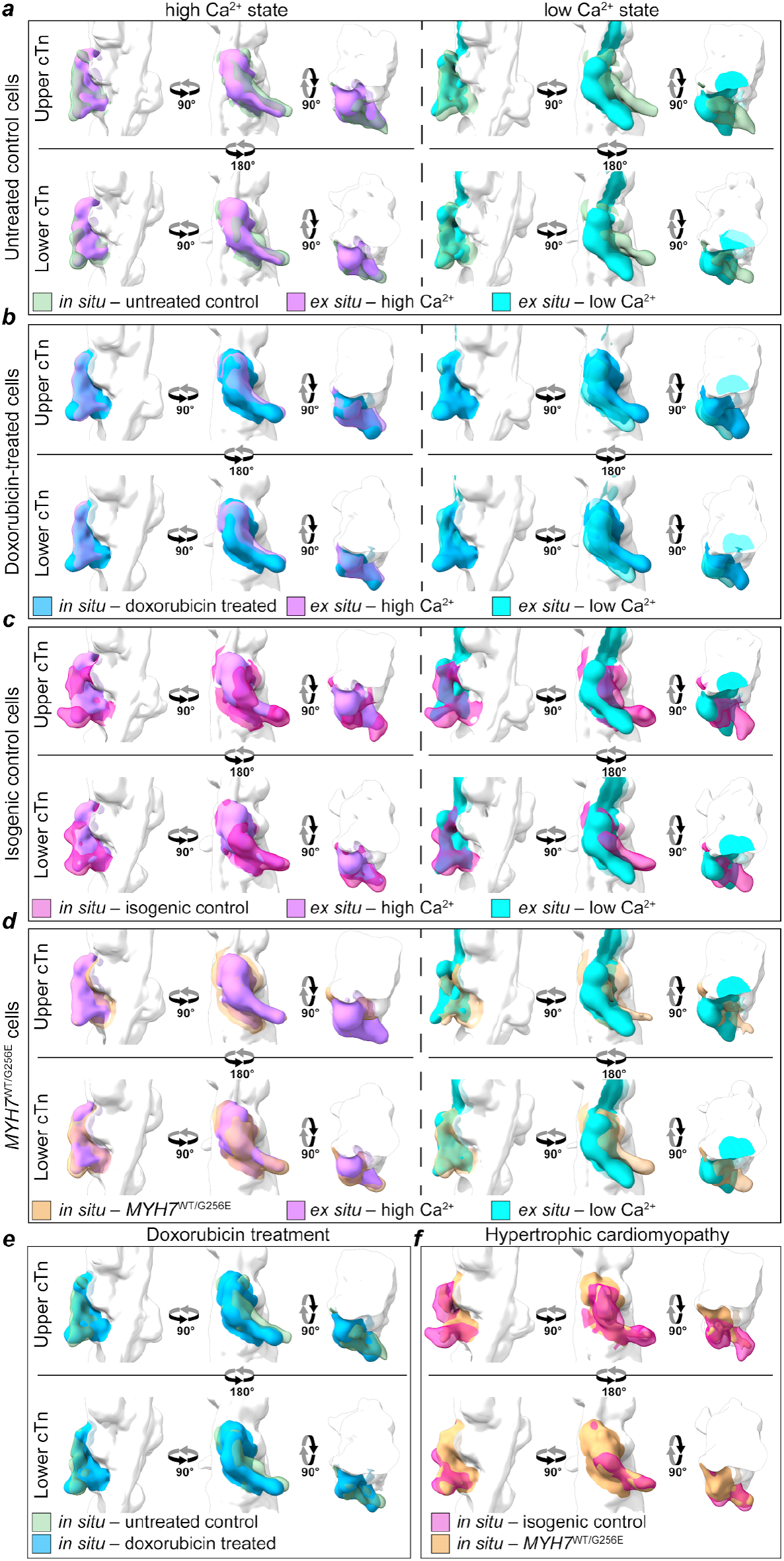
*In situ* structures of troponin complexes. Three views of the upper and lower troponin complexes (top and bottom, respectively) for pairwise comparisons of reconstructions from different sources. **(*a*)** The *in situ* reconstruction from untreated control cells (transparent green) compared to the *ex situ* high Ca^2+^ state (purple, left, EMD-0729) and the *ex situ* low Ca^2+^ state (cyan, right, EMD-0728). **(*b*)** The *in situ* reconstruction from doxorubicin-treated cells (blue) compared to the *ex situ* high Ca^2+^ state (transparent purple, left) and the *ex situ* low Ca^2+^ state (transparent cyan, right). **(*c*)** The *in situ* reconstruction from isogenic control cells (transparent hot pink) compared to the *ex situ* high Ca^2+^ state (purple, left) and the *ex situ* low Ca^2+^ state (cyan, right). **(*d*)** The *in situ* reconstruction from *MYH7*^WT/G256E^ cells (transparent gold) compared to the *ex situ* high Ca^2+^ state (purple, left) and the *ex situ* low Ca^2+^ state (cyan, right). **(*e*)** The *in situ* reconstruction from untreated control cells (transparent green) compared to doxorubicin-treated cells (blue). **(*f*)** The *in situ* reconstruction from isogenic control cells (transparent hot pink) compared to *MYH7*^WT/G256E^ cells (gold).

**Figure S6:**
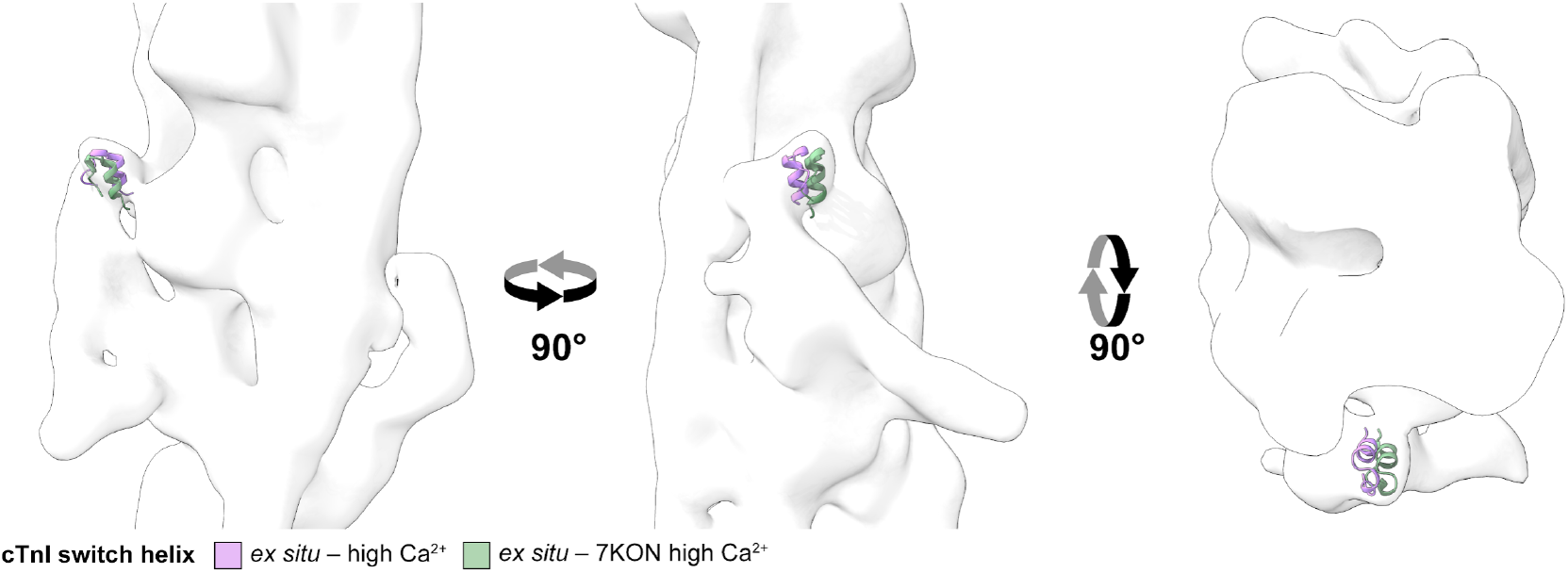
A protrusion at the cTnI switch helix in the reconstruction of the upper troponin complex from untreated control cells. Three views of the reconstruction of the upper troponin complex from untreated control cells. A density protrusion at the cTnI switch helix is better fit by a model of troponin in a high Ca^2+^ state from Risi et al^56^. (green, 7KON) than the high Ca^2+^ *ex situ* model (purple, 6KN8).

## Supplementary Tables

**Table S1.** Cryo-ET data collection and refinement statistics of vitrified and milled hiPSC-cardiomyocytes.

|  | Thin filament from isogenic control (*MYH7*<sup>WT/WT</sup>) cells |  | Thin filament from *MYH7*<sup>WT/G256E</sup> cells |  | Thin filament from untreated hiPSC-CMs |  | Thin filament from doxorubicin-treated cells |  |
| --- | --- | --- | --- | --- | --- | --- | --- | --- |
| Microscopy |  |  |  |  |  |  |  |  |
| Microscope | Titan Krios G3i |  | Titan Krios G3 |  | Titan Krios G3i CFEG |  | Titan Krios G3i |  |
| Voltage (kV) | 300 |  | 300 |  | 300 |  | 300 |  |
| Camera | K3/BioQuantum, CDS |  | K3/BioQuantum |  | K3/BioQuantum, CDS |  | K3/BioQuantum, CDS |  |
| Slit width (eV) | 15 |  | 20 |  | 15 |  | 15 |  |
| Pixel size (Å) | 2.13 |  | 2.20 |  | 2.13 |  | 2.13 |  |
| Defocus range (µm) | 3.5-5.0 |  | 3.5-5.0 |  | 3.5-5.0 |  | 3.5-5.0 |  |
| Tilt range<sup>a</sup> (pre-tilt, increment) | -48°/+48° (-9°, 3°) |  | -60°/+60° (0°, 2°) |  | -48°/+48° (-9°, 3°) |  | -60°/+60° (0°, 3°) |  |
| Tilt scheme | Dose-symmetric |  | Dose-symmetric |  | Dose-symmetric |  | Dose-symmetric |  |
| Total dose (e-/Å<sup>2</sup>) | ~105 |  | ~120 |  | ~105 |  | ~115 |  |
| Number of cells/lamellae | 3 |  | 3 |  | 3 |  | 3 |  |
| Number of tomograms | 12 |  | 8 |  | 10 |  | 14 |  |
| 3D refinement statistics | Without the troponin complex | With the troponin complex | Without the troponin complex | With the troponin complex | Without the troponin complex | With the troponin complex | Without the troponin complex | With the troponin complex |
| EMDB # | 49344 | 49349 | 49346 | 49351 | 49345 | 49350 | 49348 | 49352 |
| Number of sub-volumes | 16,451 | 1,463 | 15,147 | 2,114 | 9,851 | 721 | 7,555 | 2,737 |
| Symmetry | C1 | C1 | C1 | C1 | C1 | C1 | C1 | C1 |
| Resolution (Å) | 8.7 | 19.5 | 9.3 | 12.4 | 9.1 | 18.0 | 9.5 | 12.7 |

<sup>a</sup> Tilt angle range used only for data collection relative to the pre-tilt.

## Supplementary Digital Files

**Digital Files S1:** A digital pattern file of rectangles with a 1:7 aspect ratio and areas of 1500 μm^2^ (14.64 μm by 102.465 μm), specifically designed for micropatterning electron microscopy grids for cardiomyocytes.

**Digital Files S2:** A digital pattern file of rectangles with a 1:7 aspect ratio and areas of 2000 μm^2^ (16.903 μm by 118.32 μm), specifically designed for micropatterning electron microscopy grids for cardiomyocytes.

